# Abnormal multisensory facilitation patterns relate to disorganized thinking severity and cognitive decline in schizophrenia

**DOI:** 10.1101/2024.07.26.605377

**Authors:** Kimberly K Rogge-Obando, Brian A Coffman, Julia M Stephen

**Author notes:** **Corresponding Author:** Julia M Stephen, The Mind Research Network, 1101 Yale Blvd NE, Albuquerque, NM 87106. Co-Author: Kimberly K Rogge-Obando. Co-Author: Brian A Coffman.

## Abstract

Past research has demonstrated that patients with schizophrenia (SP) have visual processing and multisensory integration deficits. Additional studies report that sensory abnormalities are related to positive symptoms. To further understand how multisensory abnormalities relate to positive symptoms, we administered a multisensory integration task requiring the evaluation of perceived distance from auditory, visual, and multisensory stimuli with varying synchrony as well as clinical and neurocognitive assessments. Overall, patients had greater facilitation than healthy controls and the near synchronous condition had the most facilitation in comparison to other conditions. To further examine how multisensory facilitation relates to symptom severity, we performed a Ward cluster analysis that grouped participants by their multisensory facilitation profile. In contrast to what was expected, none of the Ward clusters were populated by a single group. Patients in cluster 3 had a significantly greater disorganization factor score than those in cluster 1. Our in-depth comparison between Ward clusters and neuropsychological tests reveal patients with greater multisensory facilitation experience the most cognitive deficits. Overall, our results demonstrate that multisensory integration is related to behavioral and cognitive deficits in complex ways. Further research is needed to understand the relationship between multisensory integration and schizophrenia symptomology.

**Highlights:**

- Patients with schizophrenia have greater multisensory facilitation in perceived synchronous conditions
- Multisensory facilitation patterns are heterogenous across patients with Schizophrenia.
- Different multisensory patterns relate to cognitive decline and disorganized thinking in Schizophrenia.

## 1. Introduction

Individuals living with schizophrenia (SP) experience positive (delusions, hallucinations and disorganized thinking) and negative (blunted affect, and social withdrawl) symptoms that interfere with daily functioning and quality of life (Evensen et al., 2016; Kay et al., 1967; Sullivan et al., 2019). While positive and negative symptoms have been thoroughly investigated, many studies continue to indicate patients experience basic sensory abnormalities (Gröhn et al., 2022). Additionally, some studies found that sensory abnormalities had a direct relationship to symptom severity. (Dalal, 2019; Gröhn et al., 2022; Kaufmann et al., 2015).

Unisensory and multisensory studies demonstrate SP have impairments in their visual and auditory systems (Lalor et al., 2012; Martínez et al., 2013, 2019). Butler et al., (2005) compared electroencephalography (EEG) data between SP and healthy control (HC) in the magnocellular pathway. When the stimulus was reduced to amplify signals in the magnocellular pathway, SP had an attenuated amplitude of steady-state visual evoked potentials. Source analysis revealed the reduced amplitudes propagated into dorsal and ventral streams (Butler et al., n.d., 2007). Similarly, our prior multisensory EEG study (Stone et al. 2011) identified a significant reduction in visual response amplitude in occipital channels and auditory N100 amplitude over auditory cortex in patients with schizophrenia (SP) relative to HC. (Stone et al. 2011). However, N100 amplitude was less impaired in responses to multisensory stimuli, suggesting that SP have deficits in unisensory responses that are resolved with multisensory (AV) stimuli (Stone et al., 2014). Our most recent study (Sanfratello et al., 2018) revealed less activation in the dorsal visual stream and a delayed peak activation relative to controls using a similar version of an auditory/visual multisensory task. Where the reduced response amplitude in parietal cortex was related to greater symptoms in SP. These results indicate a specific dorsal stream deficit in SP which may influence multisensory integration and is related to SP symptomology.

Different multisensory studies have related facilitation to SP symptomology. (Williams et al., 2010) performed RT quantile analysis revealing SP had less multisensory benefit in comparison to HC. HC maintained multisensory facilitation up to the 80^th^ percentile whereas SP only maintained multisensory facilitation of reaction time until the 50^th^ percentile. Increased symptom severity was associated with less multisensory benefit. Furthermore, patients who experienced both auditory and visual hallucinations had a significant decrease of multisensory facilitation relative to participants who experienced only one type of sensory (auditory or visual) hallucination.

An explanation for a relationship between multisensory integration and positive symptomology might be related to the simultaneity window, a time window within which asynchronous stimuli are perceived as synchronous. (Foucher et al., 2007) employed a parametric multisensory task that incrementally varied the delay between an auditory and visual stimulus by 10 ms up to a maximum offset of 150 ms and asked participants to determine if the visual stimulus was synchronous with the auditory stimulus. They found SP perceived auditory and visual stimuli as synchronous at larger interstimulus offsets than HC. Similarly, (Zvyagintsev et al., 2017) determined SP had significantly faster response times in comparison to HC in a streaming bouncing illusion with auditory/visual offsets and abnormal binding between auditory and visual cues. The broader simultaneity window has been theorized to contribute to multisensory deficits because the larger the window the higher the probability of inaccurately associating unrelated events. Inaccurate associations between stimuli have been speculated to promote positive symptoms in SP (Dalal, 2019). In contrast, (Stevenson et al., 2017) found a significant correlation between the temporal binding window and hallucination severity, where decreased severity was associated with an increased, or “leaky” temporal binding window and multisensory integration of temporally disparate events. These conflicting results emphasize the need to understand the relationship between multisensory integration, temporal binding and SP symptomology.

To summarize, many studies demonstrate that SP have central-peripheral visual deficits, multisensory deficits, and temporal binding abnormalities. Interestingly, many studies link multisensory integration deficits to positive symptoms; however, little is known about how heterogeneity in multisensory integration relates to symptomology. In this study, we administered an auditory-visual multisensory integration task that manipulated visual field (central/peripheral), loudness of auditory stimuli (loud/quiet), and synchrony (synchronous/asynchronous), and related the results to neuropsychological outcomes and symptomology. From Williams et al. (2010), we hypothesized that SP would have a significant decrease of multisensory facilitation relative to HC, and furthermore that these deficits would be associated with greater positive symptoms. Based on dorsal stream deficits, our second hypothesis was multisensory RT facilitation would be reduced for the peripheral vs central visual stimulus in SP vs. HC. Lastly, we hypothesized that SP would reveal more facilitation in the asynchronous condition relative to HC due to the wider window of integration in SP. Based on the Research Domain Criteria (RDoC) approach to further examine subgroups within the schizophrenia spectrum, we performed an exploratory Ward cluster analysis to identify the characteristics of individuals with similar multisensory response patterns (Cuthbert, 2015). Afterwards, we examined cluster-level differences in positive symptoms and disorganization. Finally, a post-hoc analysis of neuropsychological differences between the Ward cluster groups was performed to explore differences in cognitive abilities between clusters. Stone et al. (2014) and Sanfratello et al. (2018) briefly report the multisensory behavioral results described here but focused on the neurophysiological effects of multisensory stimulation, with the current analysis focusing on the relationship between behavioral measures of multisensory facilitation, symptomology and cognitive functioning.

## 2. Methods

### 2.1. Participants

The study was approved by the University of New Mexico Health Sciences Center Human Research Review Committee and complied with the Declaration of Helsinki. All participants underwent written informed consent prior to study procedures. As a part of a larger study (Stone et al. 2014, Sanfratello et al. 2018) we obtained behavioral measures in 63 schizophrenia patients (SP) and 67 age-matched healthy control (HC) participants. Participant characteristics are presented in Table 1. All participants were free of neurological disorders (e.g. epilepsy) determined by a neurological exam and review of symptoms. Participants also had no history of significant head trauma (<5 minutes loss of consciousness) and no current diagnosis of substance abuse (excluding nicotine). The healthy controls had no history of psychiatric disorder (assessed with SCID-NP) and no first-degree relatives with a psychotic disorder. The SP were confirmed to have a DSM-IV diagnosis of schizophrenia or schizoaffective disorder with DSM IV SCID-IP. All SP were clinically stable with no recent medication change within 1 month of study enrollment and no change of medication across the data collection period (neuropsychological testing and behavioral testing were performed at different visits). The SP symptoms [Positive and Negative Syndrome Scale - PANSS], social functioning [University of California Performance Skills Assessment - UPSA (Mausbach et al., 2007) and antipsychotic medication dose [as olanzapine equivalents; (Gardner et al., 2010)] as well as time from first episode, time from first hospitalization and other demographic information were obtained.

**Table 1.**
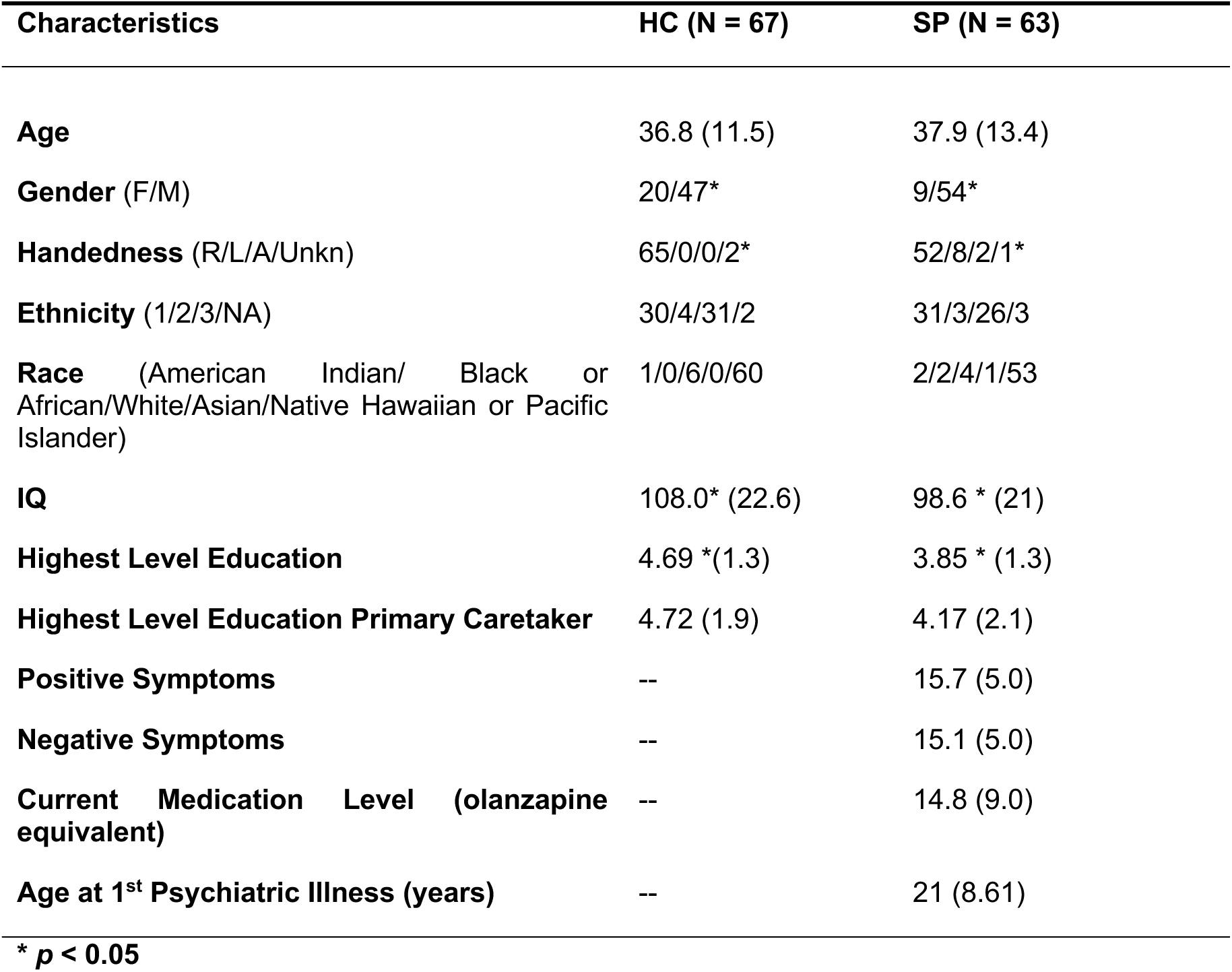
Demographics and Symptom Profile.

### 2.2. Neuropsychological Testing

All participants completed an extensive neuropsychological testing battery characterizing IQ (Wechsler Abbreviated Scale of Intelligence), premorbid IQ [Wechsler test of adult reading – WTAR (Wechsler, 2001)], and the Measurement and Treatment Research to Improve Cognition in Schizophrenia–MATRICS (Nuechterlein et al., 2008) and the Brief Assessment of Cognition in Schizophrenia (BACS) (Keefe et al., 2006) cognitive assessments.

### 2.3. Behavioral Task

During MEG measurements (Sanfratello et al., 2018), participants performed an auditory/visual multisensory integration task. The task (Stephen et al. 2010) was a discrimination choice reaction time task with visual and auditory stimuli (Lovelace et al., 2003). These stimuli were presented in a soccer field background (see Fig. 1) to provide context for the stimuli. The visual (V) stimulus (soccer ball) was presented at one of two positions (centered at 1.8° and 8° below fixation for “far” and “near” stimuli, respectively). Participants were required to fixate on the goalie in the image, which was centered horizontally and vertically with each participant’s nasion at a distance of 1 meter. The Far and Near stimuli were scaled (stimuli subtended 1° and 2.7° visual angle, respectively) to conform with the cortical magnification factor (Virsu & Rovamo, 1979) to activate an equivalent patch of cortex across the two stimuli. The auditory (A) stimulus was a 550 Hz tone (200ms duration with 30 ms Hanning ramp up/down) presented at two different volumes (45 dB (Far) and 63 dB (Near) above hearing threshold). Hearing threshold was determined independently for each ear for each participant prior to beginning the task, which was designed to eliminate perceived volume differences across subjects due to poor ear insert placement or peripheral hearing deficits. To determine the hearing threshold, 1000 Hz tones were randomly presented above and below hearing threshold in a stepwise fashion. The participant was required to press a button whenever they heard a tone.

**Figure 1.**
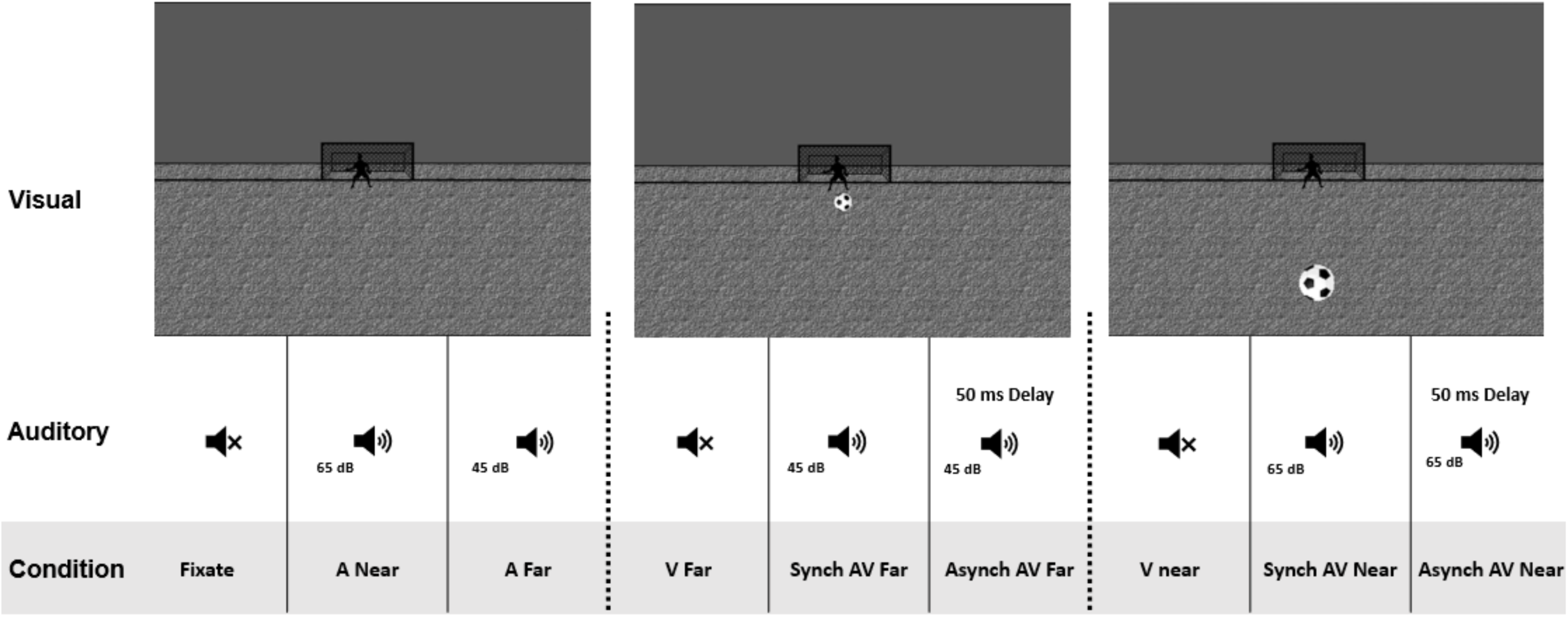
Soccer Field Paradigm Multisensory Task. Figure of the experimental conditions. The visual representation (top) is what the participant saw on the screen and the auditory sound (bottom) played for each condition.

Eight conditions were presented to the participants during the task: A Near/Far, V Near/Far, synchronous (Synch) AV Near/Far and asynchronous (Asynch) AV Near/Far (A delayed by 50 ms relative to V stimulus) see Fig. 1. The asynchronous condition emulated the natural delay between an auditory and visual stimulus which originated approximately 55 feet from the participant due to the differential speed of sound and light. A and V stimuli were congruent during AV conditions; that is, the auditory Near (Far) stimuli were always paired with visual Near (Far) stimuli. All eight conditions were presented randomly with an inter-stimulus interval (ISI) of 1500-1900 ms. The stimuli were presented in 6 blocks to provide the participant with a regular break (1-2 min.) approximately every 7 minutes. The participants were each presented with ∼150 trials/condition with the task taking approximately 45 min. For all 8 conditions, the participant was instructed to decide whether the stimulus (A, V, or AV) was near or far relative to themselves by pressing a button with the index or middle finger of their right hand, respectively. The Neurobehavioral Systems’ Presentation program was used to determine hearing threshold, present the stimuli and record the behavioral data (reaction time and percent correct). During a practice run the participant was provided with feedback on correct/incorrect trials. This feedback (ball rolled into the goal and/or cheering sound was heard for correct trials, or the ball missed the goal and/or a disappointed crowd sound was heard for incorrect trials) was also provided for 20% of the trials during the MEG task to encourage compliance throughout data collection.

### 2.4. RT Analysis

Visual RTs were faster than Auditory RTs both within and across diagnostic groups. To identify multisensory facilitation of mean RTs, we subtracted the fastest mean unisensory RT (visual) from the multisensory mean RT (Laurienti et al., 2006). Although mean RT facilitation provides a measure of multisensory integration, the RACE model determines if the experimental RTs are faster than the statistical facilitation predicted from the receiving input from two independent channels (auditory/visual). We performed an analysis similar to that reported by Williams et al. (2010) based on the Miller Race model (Ulrich et al., 2007). We modified the output to be consistent with the results presented by (Molholm et al., 2002) by comparing 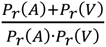 to allow us to look at facilitation across the entire quantile curve. This allowed us to better understand the relationship between the quantile facilitation and the mean RT facilitation.

### 2.5. Statistical Analysis

Statistical analyses were performed using IBM SPSS version 20. RTs and were compared statistically using a repeated measures analysis of variance (RM-ANOVA). Results were compared with condition (A, V, AV), location (Near/Far) and synchrony (Synch vs. Asynch) as within subjects’ factors and group (SP vs. HC). To further understand the results, we performed a Ward cluster analysis. We decided to use the 5^th^ and 90^th^ quantile because this models the participants quickest, and slowest response time. Additionally, we used the near synchronous condition as a variable because this condition had the greatest amount of multisensory integration. The Ward cluster method identified 3 clusters. We observed each cluster result and ensured each was distinct (see Figure 3A). The patient symptom factors and medications were then compared across the 3 clusters using one-way ANOVA. Furthermore, the neuropsychological measures were compared across clusters to determine if cognitive abilities were related to specific groups based on multisensory facilitation variables. Lastly, R-studio was used to conduct a Kruskal-Wallis test for the positive and disorganized factors followed with a post-hoc Dunnett’s test.

## 3. Results

Demographic information is presented in Table 1. As expected, there was a significant difference in IQ between SP and HC and a significant difference in the highest level of education. However, there was no significant difference in highest level of education of the primary caretakers to age 18. The groups were matched (p > 0.05) on age, race, ethnicity but not on gender or handedness.

The results are presented for both the mean RT and the quantile RTs throughout. The facilitation of the mean RT was assessed by calculating the difference between the multisensory (AV) RT and the visual RT (the fastest unisensory RT for both groups). This facilitation was compared using a 3-way mixed measures ANOVA with distance and synchrony as within subject factors and diagnosis as the between subject factor. We found a main effect of distance (F(1,128) = 209.3, p < 0.001) and synchrony (F(1,128) = 6.4, p = 0.013) with no significant two- or three-way interactions. Peripheral (near) stimulation had greater facilitation relative to central (far) visual stimulation and synchronous AV condition showed greater facilitation than the asynchronous condition (A delayed by 50 ms relative to V). Finally, there was a significant group difference (F(1,128) = 13.85, p = 0.0003) with SP having greater facilitation than HC.(Figure 2)

**Figure 2.**
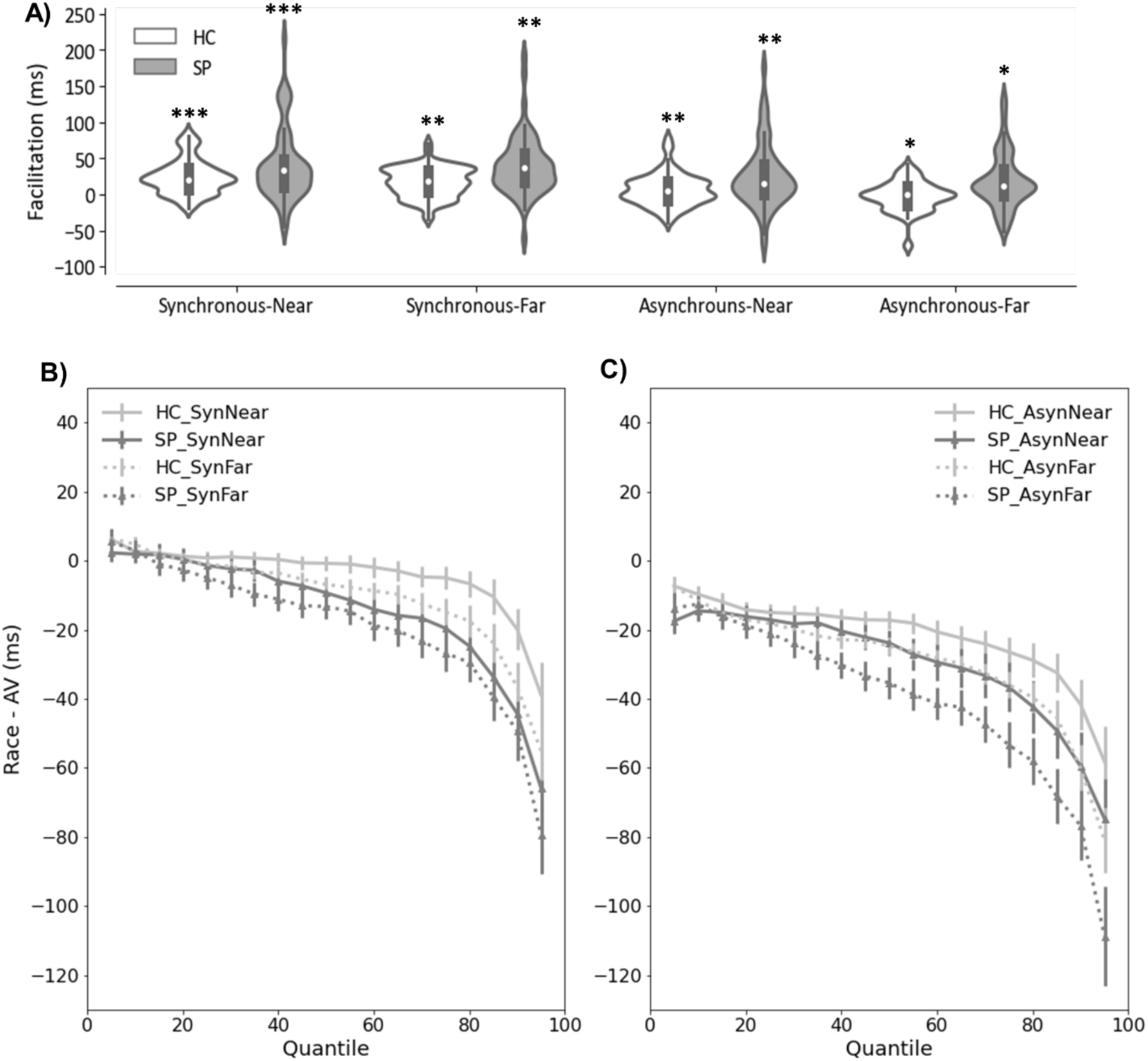
The RT Mean Facilitation and Quantile Results of Facilitation for Conditions. A) Is a violin plot for the RT Mean Facilitation of each condition between groups where a * represents significance of group, ** represents significance of synchronous or near condition. B) Is a linear plot for RT quantile synchronous results between SP and HC.

Similarly, we calculated the difference between the Miller RACE model RTs and experimental AV RTs at each quantile. These values were compared statistically using a 4-way mixed measures ANOVA with distance, synchrony and quantile as within subject factors and diagnosis as the between subject factor. Again there was a group difference (*F*(1,128) = 4.26, *p* = 0.041) with SP (25.7 sem 3.3) showing greater facilitation (AV RTs were faster than the Miller race model) across quantiles relative to HC (16.4, 3.2). Similar to the mean RTs there were significant main effects for quantile (F(1.6, 201.7) = 84.9, p < 0.001), synchrony (F(1,128) = 222.0, p < 0.001) and distance (F(1,128) = 18.1, p < 0.001). There was also a significant interaction (F(2.6,329.5) = 7.7, p < 0.001) between quantile and distance. In general, the amount of facilitation increased by quantile. The facilitation was greater for synchronous (30.3 SEM 2.5) versus asynchronous (11.8, 2.2) and greater for near (24.7, 2.4 - peripheral visual stimulus) than for far (17.4, 2.4 - central visual stimulus). (Figure 2)

To better understand the observed variability in facilitation across participants we performed a Ward cluster analysis using three variables, the synchronous near 5^th^ and 90^th^ quantile facilitation value and the mean RT facilitation value. These were selected based on the pattern that revealed the greatest group differences (Figure 2A). Using the Ward clustering method, we identified 3 distinct facilitation clusters (Figure 3A). Two of the clusters were dominated by SP and one cluster was dominated by HC participants, but none of the clusters were populated by only one group.

**Figure 3.**
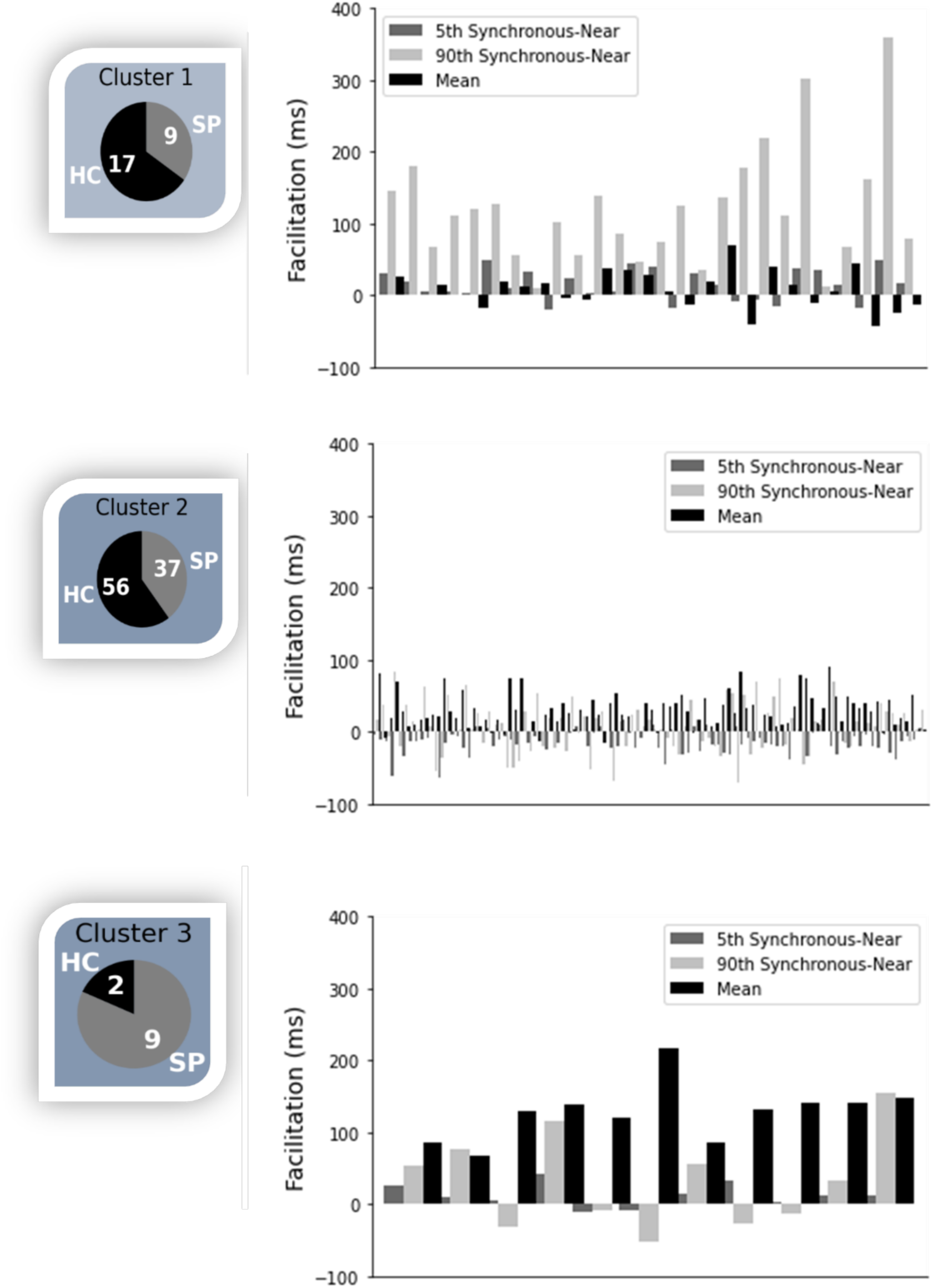
Ward Cluster Multisensory Results Distribution. Multisensory characteristics within clusters and group membership distribution. Where the black/ grey in the pie chart represents the number of HC/SP in each cluster.

**Figure 4.**
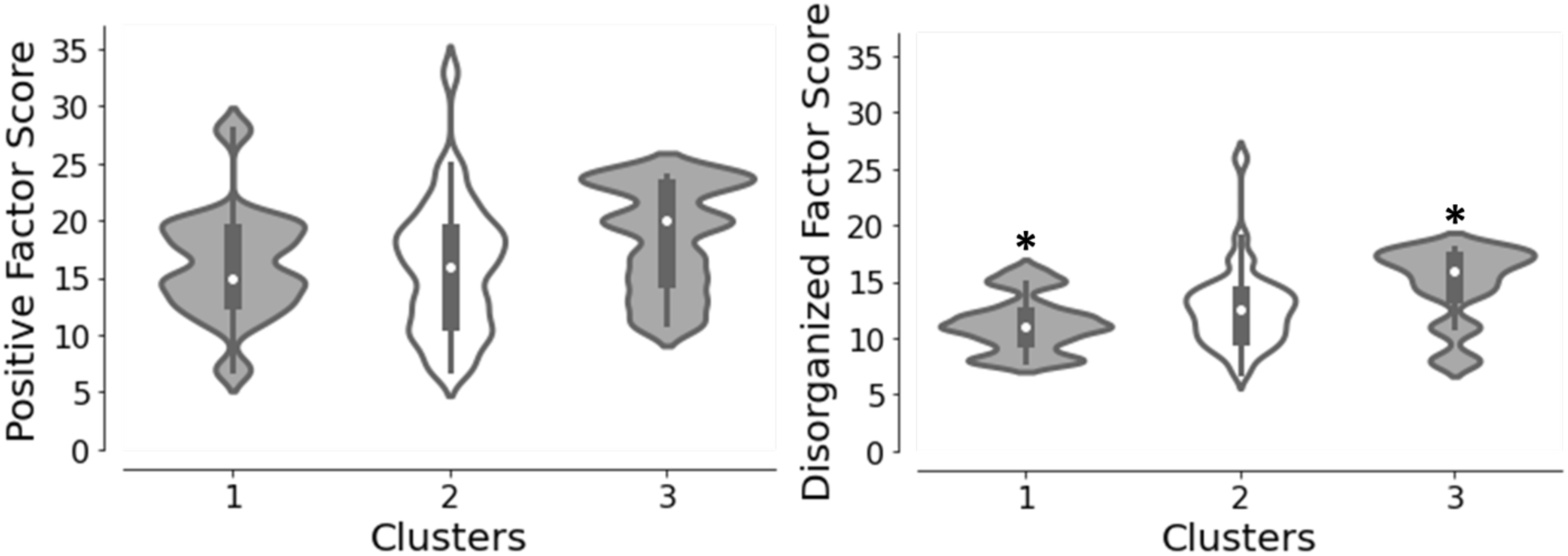
Ward Cluster Symptom Factor Comparison. Violin plots of the positive symptom score disorganized thinking score from the PANSS by Ward Cluster.

We performed a general linear model analysis of the neuropsychological test results to examine differences by Ward cluster. Overall, the results were consistent with individuals in cluster 3 generally performing worse than individuals in clusters 1 & 2 and overall individuals from cluster 1 performed best although generally clusters 1 & 2 did not differ significantly from each other (Table 2).

**Table 2.** Comparison of Cognitive Assessment scores by Ward Cluster Grouping.

| Assessment | F-value (2, 106) | p-value | Post-hoc results |
| --- | --- | --- | --- |
| WTAR_StandardScore | 5.5 | 0.005 | 1,2 > 3<br>2 > 3 |
| WTAR_Difference | 6.6 | 0.002 | 1,2 > 3 |
| WASI_Vocab_TScore | 2.5 | 0.086 |  |
| WASI_Similarities_TScore | 5.8 | 0.004 | 2 > 3<br>n.s. |
| WASI_Verbal_TScore | 4.5 | 0.013 | n.s. |
| WASI_Verbal_IQ | 4.3 | 0.015 | n.s. |
| WASI_BlockDesign_TScore | 14.2 | < 0.001 | 1,2 > 3 |
| WASI_MatrixReasoning_TScore | 11.2 | < 0.001 | 1,2 > 3 |
| WASI_Performance_TScore | 16.7 | < 0.001 | 1,2 > 3 |
| WASI_Performance_IQ | 14.7 | < 0.001 | 1,2 > 3 |
| WASI_Sum_of_TScores | 7.8 | 0.001 | 2 > 3 |
| WASI_Sum_IQ | 1.8 | 0.17 |  |
| WAIS_SymbolSearch_ScaleScore | 4.7 | 0.011 | 2 > 3 |
| WAIS_Coding_ScaleScore | 9.2 | < 0.001 | 2 > 3 |
| WAIS_PSI | 1.3 | 0.275 |  |
| TMT_A_TScore | 15.5 | < 0.001 | 1,2 > 3 |
| BACS_SC_TScore | 12.9 | < 0.001 | 1,2 > 3<br>2 > 3 |
| HVLT_Trials_TScore | 2.5 | 0.085 |  |
| HVLT_Delay_TScore | 4.7 | 0.011 | 2 > 3 |
| HVLT_DI_TScore | 2.0 | 0.14 |  |
| WMSIII_SS_TScore | 7.9 | 0.001 | 2 > 3 |
| **LNS_TScore** | 5.5 | 0.005 | 1,2 > 3 |
| **NAB_Mazes_TScore** | 11.0 | < 0.001 | 1,2 > 3 |
| **BVMT_Trials_TScore** | 3.3 | 0.042 | n.s. |
| **BVMT_Trials_Delay_TScore** | 2.4 | 0.094 |  |
| **FAS_Sum_TScore** | 4.4 | 0.015 | 2 > 3 |
| **Animal_Fluency_TScore** | 3.2 | 0.045 | n.s. |
| **MSCEIT_TScore** | 2.5 | 0.086 |  |
| **CPTIP_TScore** | 7.2 | 0.001 | 2 > 3 |
| **MatricsDomain_ProcessingSpeed_TScore** | 13.1 | < 0.001 | 1,2 > 3 |
| **MatricsDomain_AttentionVigilance_TScore** | 15.6 | < 0.001 | 2 > 1,3 |
| **MatricsDomain_WorkingMemory_TScore** | 8.9 | < 0.001 | 1,2 > 3 |
| **MatricsDomain_VerbalLearning_TScore** | 2.5 | 0.082 |  |
| **MatricsDomain_VisualLearning_TScore** | 3.3 | 0.04 | n.s. |
| **MatricsDomain_ReasoningProblemSolving_TScore** | 10.9 | < 0.001 | 1,2 > 3 |
| **MatricsDomain_SocialCognition_TScore** | 2.5 | 0.088 |  |
| **MatricsDomain_OverallCompositeScore_TScore** | 12.9 | < 0.001 | 1,2 > 3 |

Where WTAR is Wechsler Test of Adult Reading, WASI is Wechsler Abbreviated Scale Intelligence, TMT A is the Trail Making Test for Attention, BACS is the Brief Assessment of Cognition in Schizophrenia, HVL T is Hopkins Verbal Learning Test, WMS III is Wechsler Memory Scale -Third Edition, LNS is Letter Number Sequencing, NAB Mazes is Neuropsychological Assessment Battery, BVMT Brief Visuospatial Memory Test-Revised, FAS is the Functional Assessment Systems, MSCEIT is The Mayer Salovey Caruso Emotional Intelligence Test, CPTIP is The Continuous Performance Test identical pairs version.

Finally, we conducted a Kruskal-Wallis test for the positive and disorganized factors reported from Emsley et al. (2002) in the PANSS symptom scores between Ward clusters. The between-subject results identified a significant difference between cluster for disorganized factor (H(2), p=0.029) but not for the positive factor (H(2), p= 0.57). The post-hoc analysis for the disorganized factor revealed a significant difference between cluster 1 and cluster 3 at p < 0.05. Patients in cluster 3 had significantly greater disorganized thinking than those in cluster 1 (Figure 3C). This finding reveals individuals with greater multisensory facilitation were also the individuals with greater cognitive deficits and disorganization. Additionally, greater multisensory facilitation in cluster 3 normalized reaction times. Unfortunately, a limitation in this exploratory investigation was the different number of individuals populating each cluster. Despite the uneven amount, each population illustrates a unique multisensory pattern and symptom profile, see supplemental Figure 3.

## 4. Discussion

There continue to be new reports of deficits in multisensory facilitation in patients with schizophrenia (SP). However, the results of the current analysis reveal a complex story. First, the patients had greater multisensory facilitation than HC for both mean RT facilitation and across quantiles relative to the Miller RACE model. Mean facilitation is the difference between the fastest unisensory response time (V) and the multisensory response time (AV). In this study, greater facilitation in the SP population originates from a faster AV response. While there is some prior evidence that patients with schizophrenia experience deficits in multisensory integration (Williams et al. 2010), other studies make it clear that experimental conditions and symptom severity complicate this narrative (Wynn et al., 2013). Wynn et al., (2014) performed a simple auditory and visual multisensory task. Their results suggested that SP and HC did not differ in multisensory facilitation, either behaviorally or in their event related potential (ERP) response. Instead, HC maintained a multisensory benefit to the 68^th^ percentile and the SP group maintained a benefit until the 58^th^ percentile. One important difference between the Williams paper and the current report is that the Williams task presented simple stimuli that required participants to press the button as quickly as possible whenever an auditory or visual stimulus was presented (i.e., reaction time). In contrast, the soccer ball task, presented high contrast sensory stimuli and required participants to perform a discrimination task rather than a simple detection task (i.e., response time). This additional complexity engages more distributed and higher-level brain networks and may explain the conflicting results. Furthermore, the facilitation in SP is largely seen near the mean, suggesting an overall facilitation of RT and explaining the mean facilitation difference by group.

Second, we hypothesized that facilitation would be greater for the central visual condition due to the known deficits in the dorsal visual stream in SP. However, greater multisensory facilitation was seen in the peripheral (near) condition. This may be better explained by attention capture than by the dorsal stream deficit. One possible explanation for the increased facilitation in patients with SP relative to HC in this study is that patients unisensory responses may be inherently noisy (Cheng et al., 2015; Freedman et al., 1987; Molina et al., 2016). This is consistent with the high noise theory of schizophrenia (Kaufmann et al., 2015). When multisensory stimuli are presented to SP, the noisy unisensory response may be enhanced by the presence of the multisensory stimulus enabling a faster reaction time. This is also consistent with the inverse effectiveness rule of multisensory integration (Stein and Meredith, 1993; Holmes, 2007). In controls, multisensory facilitation occurs more commonly with difficult to perceive stimuli – or stimuli near the perceptual threshold. In this study, the stimuli were high contrast and were not expected to be difficult to perceive for either group. Yet, if patients with schizophrenia have more internal neuronal variability or noise, the stimuli may be perceived as lower contrast relative to controls (Faisal et al., 2009). Additionally, attention has been demonstrated to impact multisensory facilitation (Talsma et al., 2010). Stimuli that are perceived to be closer to an individual are inherently more salient (Pani et al., 2018; Reynolds et al., 2003). Patients with schizophrenia are also recognized to have deficits in attention and therefore having more salient stimuli may better capture attention (Chen et al., 2000; Carter et al., 2010; Harris et al., 2007).

The mean facilitation results for synchronicity were consistent with the prior literature. Prior studies demonstrate greater multisensory facilitation happens when stimuli are synchronous, and facilitation decreases when sensory stimuli are perceived as separate events. For example, no facilitation was observed when the time between stimuli was greater than 100 ms (Foucher et al., 2007). The greater mean facilitation for the synchronous condition and the quantile results confirmed our hypothesis. Again, facilitation at RT by quantile is the difference between the Miller Race model and the multisensory RTs. Therefore, positive values of facilitation represent a faster RT experimental response than the Miller Race models predicted.

The Ward cluster analysis was performed to understand the variability in the multisensory reaction time data reported across groups. The primary question was whether variability within the schizophrenia population was contributing to the discrepant results. This was further motivated by the research domain criteria (RDoCs) framework to better understand the variability observed in clinical populations (Insel et al 2012). The Ward cluster analysis demonstrates again that there is considerable variability in multisensory facilitation in both groups (HC & SP). As discussed previously, Williams et al. (2010) indicated that the type of hallucination was related to the deficit in multisensory facilitation in SP patients. The results from this study revealed cluster 3 had the greatest multisensory facilitation and experienced the more severe symptom profile (disorganized thinking) and the poorest performance on neuropsychological tests. Interestingly, increased multisensory facilitation in this group tended to normalize their RT (the facilitation brought their delayed RTs closer to the mean RT of the other groups for multisensory conditions). This result alone suggests that patients with schizophrenia with more severe disorganized thinking may be the ones who would benefit from multisensory stimuli during intervention or cognitive training.

## 5. Conclusion

To conclude, many studies support visual deficits in SP that may affect multisensory processing and SP symptomology. The current study investigated multisensory facilitation and identified significant differences between group, distance, and synchrony. Specifically, SP in near and synchronous conditions had the greatest mean facilitation. Facilitation relative to the Miller Race model found similar significance except asynchronous conditions had greater facilitation. The Ward cluster analysis revealed that SP with the most facilitation were also those individuals with greater symptoms and poorer cognitive performance. Future studies are encouraged to investigate how symptom severity changes with multisensory facilitation to identify if an increase of multisensory facilitation leads to greater positive symptoms scores and lower cognitive scores.

## Notes

### Competing Interest Statement

The authors have declared no competing interest.

